# Pooled screening for biochemically-defined phenotypes using CRISPuRe-Seq

**DOI:** 10.1101/2023.11.15.567224

**Authors:** David T. Harris, Calvin H. Jan

**Author notes:** Electronic address.

## Abstract

Phenotypic genetic screening is a powerful approach for biological discovery. CRISPR technologies have enabled routine genome-scale pooled perturbations in cultured cells, creating an opportunity to develop screening methods capable of accessing molecularly meaningful phenotypes. Here we present a generalized strategy, named CRISPuRe-seq, that enables pooled screening of a broad range of protein-based attributes. This approach utilizes the purification of an RNA-barcoded protein-of-interest to identify causative genetic perturbations. As a proof of concept, we screened for modifiers of Interferon alpha signaling and were able to identify all of the primary components of the type 1 Interferon JAK/STAT signaling pathway.

## Introduction

CRISPR-based systems have become widely adopted as the preferred platform for genetic screening at the whole-genome scale. The BioGRID ORCS database of human CRISPR screens reports over 1500 screens, yet only approximately 20 phenotypes have been queried. The vast majority of these screens are based on growth phenotypes, where pools of cells are combined and processed in parallel. As a highly integrative phenotype, growth-based screens require thoughtful engineering of conditions to probe specific biology when the objective is not related to cell fitness. High dimensional phenotypes generated by perturb-seq^1,2^ or microscopy-based^3^ screens can provide holistic representations of perturbed states. More focused screens have been devised that utilize a fluorescent reporter to convert their biology-of-interest into a suitable fluorescent proxy that can be measured by Fluorescence-Activated Cell Sorting (FACS), but these screens can be labor-intensive and equipment-limited. Recently, this focused tool set has been expanded to include the use of barcoded RNA to monitor gene expression, as in CiBER-Seq^4^, and the use of peptide barcodes to monitor protein levels by FACS or mass cytometry, as with Pro-Codes^5^. We sought to develop a screening technique that ties the state of a protein (level, post-translational modification, etc.) with the ability to carry genetic information about the cell from which the protein was expressed.

Here we describe CRISPuRe-Seq, a novel platform for high-throughput, whole-genome CRISPR screening. This platform is built upon the biochemical purification of a protein-of-interest that is physically complexed to a barcoded RNA, which is in turn linked to the identity of a corresponding CRISPR perturbation. Physically coupling perturbation identity to a protein-of-interest enables the interrogation of cell-specific protein states using a pooled purification strategy.

## Materials and Methods

### CRISPuRe library construction

Each library plasmid contains the following relevant parts: 3rd generation Lentiviral expression components, an sgRNA driven by the human U6 promoter on the positive strand, a barcoded MS2 Hairpin-containing small RNA (BarRNA) driven by the human 7SK promoter on the negative strand, and a puromycin resistance gene driven by the EF-1a promoter (See sample file Nontarget_ci_MS2_7sk.gb). The BarRNA contains a single MS2 hairpin, a 25 nucleotide random barcode, and primer binding sites for reverse transcription and subsequent pcr amplification.

We generated CRISPuRe plasmid libraries by incorporating a BarRNA expression cassette into an already existing CRISPRi plasmid library (See sample file Initial_CrisprI_Library.gb). An existing library of CRISPRi plasmids was digested with BamHI and EciI to liberate a library of fragments containing the sgRNAs and a portion of the Ampicillin resistance gene. A plasmid containing a generic BarRNA expression cassette was digested with EcoRI and FspI to liberate a second fragment containing half of a complete BarRNA (minus the barcode), the puromycin resistance gene, and an overlapping piece of the Ampicillin resistance gene. A third barcode-library-containing fragment was generated using Klenow-based overlap extension of two oligonucleotide Ultramers from IDT (Integrated DNA Technologies)^6^. The barcode-containing ultramer was synthesized carrying a stretch of 25 randomly incorporated nucleotides. These three pieces were pooled together and combined using Gibson Assembly (NEB Gibson Assembly Master Mix-E2611L).

The assembled product was electroporated into MegaX electrocompetent cells (ThermoFisher-C640003) and was plated on large agar plates. Colonies from the plates were scraped and plasmid was purified (ZymoPure II Plasmid Maxiprep-Zymogen-D4203). Each library was cloned at a depth such that each protospacer is paired with approximately 30 unique barcodes. While there are many barcodes per protospacer, there are very few barcodes shared between protospacers. Once constructed, the plasmid pool was amplified by pcr and the protospacer/barcodes were deep sequenced to generate a look-up table of protospacer-barcode pairings.

### Lookup table construction

A single sublibrary containing BarRNA and sgRNA for coding genes associated with inflammation and innate immunity was cloned and sequenced using a Hiseq 4000. Raw sequence reads were demultiplexed and putative barcode/protospacer pairs were detected using string matching for the sequence surrounding each barcode and protospacer. Putative barcode/protospacer pairs with fewer than 3 sequence reads were removed. The remaining list was then compared to the list of known protospacers and only those barcode/protospacers with an identical protospacer match were kept. Barcodes that were shared between multiple protospacers were removed.

### Screen procedure

50 million K562 cells stably expressing KRAB-dCas9 and carrying an ISRE reporter were transferred to a T175 flasks containing 100 mL of RPMI 1640 + 10% FBS + Glutamax + 8 ug/ml polybrene media. Six ml of lentivirus-containing supernatant (titrated for an MOI of ∼.3) was added (AM Day 0). Cells were infected for 24 hrs. The following day, cells were spun down and virus-containing media was replaced with fresh RPMI + 10% FBS + Glutamax and cells were transferred back to a 250 ml spinner flask (AM Day 1). Puromycin selection at 1 ug/ml was initiated 24 hrs later (AM Day 2). Selection was continued for 2 days (Days 2 and 3). Cells were spun down and puromycin-containing media was removed and replaced with media without puromycin (AM Day 4). Cells were allowed to grow for 24 hrs in the absence of puromycin prior to any screening.

On treatment day (PM Day 5), 20 million library-carrying cells were split into two (10 million each) T25 flasks at .5 X10^6^ cells/ml. Cells were treated with interferon overnight. On screening day (AM Day 6), cells were spun down for 10 minutes at 500 rcf at 4 C. Cell pellets were resuspended in 1 ml of cold PBS and then spun down again. PBS was removed prior to cell lysis.

### Cell lysis for screening

10 million cells were resuspended in .5 ml of cold hypotonic lysis buffer (20 mM HEPES ph 7.9, 2 mM MgCl2, 10% glycerol, 0.1% IGEPAL, 0.5 mM DTT, 150 mM NaCl, Protease/Phosphatase Inhibitor (Halt Protease and Phosphatase Inhibitor Cocktail (Fisher Scientific-P178444) plus competing hairpin RNA (Integrated DNA Technologies) at a concentration of 10 ug per ml. Cells were left on ice for 30 minutes for complete lysis. Trituration beyond complete resuspension is not needed. Cells were then spun at 7500 rcf at 4 C for 5 minutes to pellet nuclei and cell debris.

### Total RNA isolation

150 ul of supernatant were transferred to 1 ml of Trizol (TRIzol Reagent-Life Technologies-15596018). 200 ul of chloroform was added to each 1 ml tube of Trizol/sample. Samples were vortexed for 15 seconds and incubated at room temperature for 2 to 3 minutes. Samples were transferred to Phasemaker tubes (Life Technologies-A33248) for phase separation. Phasemaker tubes were spun for 5 minutes at 12,000 rcf. The aqueous phase was transferred to a new tube. We then added 1 ul of GlycoBlue (Life Technologies-AM9516) and 500 ul of isopropanol to each tube. We then incubated the tubes at -80 for 30 minutes or until the sample was frozen. We then thawed the sample and spun it for 10 minutes at 12,000 rcf at 4 C. Supernatant was removed and the pellet was washed twice with 1 ml of 75% ethanol, briefly vortexing and spinning at 7500 rcf for 5 minutes for each wash. Ethanol was removed by pipetting and the pellet was allowed to air dry. The Total RNA pellet was resuspended in 20 ul of water.

### Small RNA enrichment

To improve the amplification efficiency of the BarRNA from the total RNA fraction, we performed a small RNA enrichment step prior to cDNA synthesis. For each sample, we performed two enrichments in parallel. 10 ug of total RNA was diluted to a volume of 100 ul in water. We then added 70 ul of SpriSelect magnetic beads (Beckman Coulter INC-B23317) and mixed by pipetting. We let this sit for 5 minutes at room temperature. We then moved the sample to a magnetic rack and allowed the beads to fully stick to the side of the tube. We then transferred the supernatant containing the enriched smaller RNAs to a new tube.

We then performed a second isopropanol precipitation on the enriched fraction by adding 600 ul of water, 1 ul of GlycoBlue and 750 ul of isopropanol. We incubated the sample at -80 for an hour and continued with the RNA precipitation as above. The final small RNA enriched fraction was resuspended in 10 ul of water.

Alternatively, small RNAs can be enriched from total RNA using SPRISelect beads (Beckman Coulter-B23317) by doing a 2-sided purification with ratios of 0.5X and 2.5X.

### Immunoprecipitation

After cell lysis, supernatant was incubated with 25 ul of M2 anti-FLAG magnetic beads (Sigma Aldrich-IPFL00005) rotating at 4 C for 90 minutes. Beads were washed (inverted three times) with 1 ml of cold wash buffer (Lysis Buffer without IGEPAL) three times and were then resuspended in 20 ul of RNAse-free water.

### cDNA synthesis for screen analysis

cDNA synthesis was performed following the Superscript II Reverse Transcriptase (ThermoFisher-18064014) (or Maxima H Minus Reverse Transcriptase-ThermoFisher-EP0752) protocol, with minor modification. For cDNA synthesis from the IP fraction, we used the resulting bead solution (including the beads) as template in a 40 ul reaction. For the Lysate fraction, we used all 10 ul of the small-RNA enriched fraction as template. The amount of added RT primer, which contained a Unique Molecular Identifier (UMI) sequence, was 5 picomoles instead of 2 picomoles per 20 ul of reaction. Following incubation at 42 C for 50 minutes, 1 ul of EXO-SAP-IT Express (Life Technologies-75001.1.ML) was added to the reaction. This reaction was incubated at 37 C for 5 minutes before being inactivated at 80 C for 5 minutes. cDNA reactions containing beads were raised to 95 C for two minutes and then quickly moved to a bar magnet. The supernatant containing the cDNA reaction was then transferred to a new tube and subsequently used for sequencing library preparation.

### Sequencing library construction

Sequencing library construction was carried out in two rounds of pcr. Round 1 amplifies the cDNA and attaches common adapters to each product. Round 2 further amplifies and adds specific indices for Illumina sequencing. 5 ul of cDNA reaction was used in each of 4 x 50 ul Round 1 PCRs per sample using Phusion polymerase in HF buffer (Life Technologies-F531L). The 4 Round 1 PCRs were then pooled and 5 ul of that mix were used as template for a 50 ul Round 2 PCR. The Round 2 PCR underwent 12 amplification cycles. Each reaction was then either SPRI purified or run on a 1.8% agarose gel and gel purified. Primers used for reverse transcription and library construction can be found in Supplemental_File_1.xls.

### Screen barcode processing

Barcode and UMI sequences were extracted from each demultiplexed fastq.gz file using a custom bash script (CRISPuRe_mageck2.sh). For each condition, barcodes were matched to their corresponding protospacer sequence and the number of unique UMI/Barcode pairs for each protospacer were counted. Unique barcode counts were then summed for each corresponding protospacer sequence. Protospacer counts from paired samples (Total RNA vs Pulldown RNA) were compiled into a count table. This count table was then processed using MAGeCK^7^ to determine which genes/perturbations were enriched for each sample. All screens were performed and analyzed in duplicate using the MAGeCK “paired” function. MAGeCK output from all screen processing can be found in Supplemental_File_1.xls.

### Protein-Barcode association

HEK 293T cells that stably expressed either MCP-mClover3-Flag(3x) (See supplemental file pCDH_CMV_CPuRe.gb) or MCP-mClover3 (See supplemental file pCDH_CMV_CPuRe_NoFlag.gb) were generated by lentiviral transduction. MCP-mClover3-Flag(3x) cells were transduced with lentivirus for the expression of a single sgRNA-barcode (target) pair. MCP-mClover cells were transduced with lentivirus for the expression of a pool of many sgRNA-barcode (non-target) pairs. Target and non-target barcode expressing cells were mixed at a ratio of 1:100 in a total of approximately 10 million cells. Cells were lysed in a hypotonic lysis buffer in the presence or absence of “Competitor RNA”, cytosolic fractions were isolated by centrifugation and FLAG immunoprecipitation was performed for 1 hour. A 50 ul aliquot of cytosolic fraction was used to quantify total “target” and “non-target” barcode levels in the lysate.

For the protein-barcode stability analysis, FLAG immunoprecipitation was performed on the cytosolic lysate of 1 million MCP-mClover3-Flag(3x) target-barcode expressing cells. After 1 hour of binding to the IP beads, the target lysate was replaced with cytosolic lysate from 10 million MCP-mClover3 non-target-barcode expressing cells. The non-target lysate and bead mixture was rotated at 4C for increasing amounts of time. At each final timepoint, the beads were washed three times and were then further processed for cDNA sequencing and subsequent analysis. All sequencing for initial Protein-Barcode associations were performed on an Illumina MiSeq.

### Flow Cytometry

For each sample 500,000 cells were spun down in a 1.5 ml tube. Each sample was washed once in PBS and then resuspended in 100 ul of PBS + 1 ul of anti-CD47 PerCP-Cy5.5 conjugated antibody. Cells were incubated at 4 degrees for 1 hr and then washed once with 1 ml of PBS. Cells were spun down and resuspended in 100 ul of PBS + 2% FBS. Cells were then analyzed on a BD Fortessa using the PerCP-CY5.5 filter.

### ISRE Reporter Cells

Either 5 or 7 copies of an IRF-ISRE element^8^ were inserted into a lentiviral expression construct upstream of a TATA-box containing minimal promoter and the MCP:mClover3:Flag(3X) reporter coding sequence (See supplemental files pCDH_ISRE_5X_CPuRe.gb and pCDH_ISRE_7X_CPuRe.gb. Lentivirus was generated by transfection into HEK293T cells and viral supernatant was subsequently used to infect CRISPRi-enabled K562 cells carrying KRAB-dCas9. Cells were selected with antibiotic and clonal populations were generated. A 7x ISRE reporter line was initially selected based on strong mClover3 fluorescence. A 5x ISRE reporter line was selected based on reduced mClover3 fluorescence relative to the 7x ISRE cell line.

## Results

### Screening by protein purification

CRISPuRe-Seq comprises two main components: a phenotypic reporter consisting of a protein-of-interest (POI) fused to the RNA-binding MS2 Coat Protein (MCP) (Figure 1A), and a perturbation library expressing sgRNAs paired with barcoded small RNAs (BarRNA) (Figure 1B). Each barcode is paired in *cis* with a co-expressed genetic perturbagen, typically a CRISPR sgRNA. The CRISPuRe-Seq phenotypic reporter is physically tethered to the genotype-identifying BarRNA through the highly specific and stable MCP/MS2 hairpin interaction^9^. In this way, the POI is physically linked to the BarRNA, and therefore carries information about the genotype of the cell that expressed it. This linkage can then be used to interrogate various properties of the POI.

**Figure 1.**
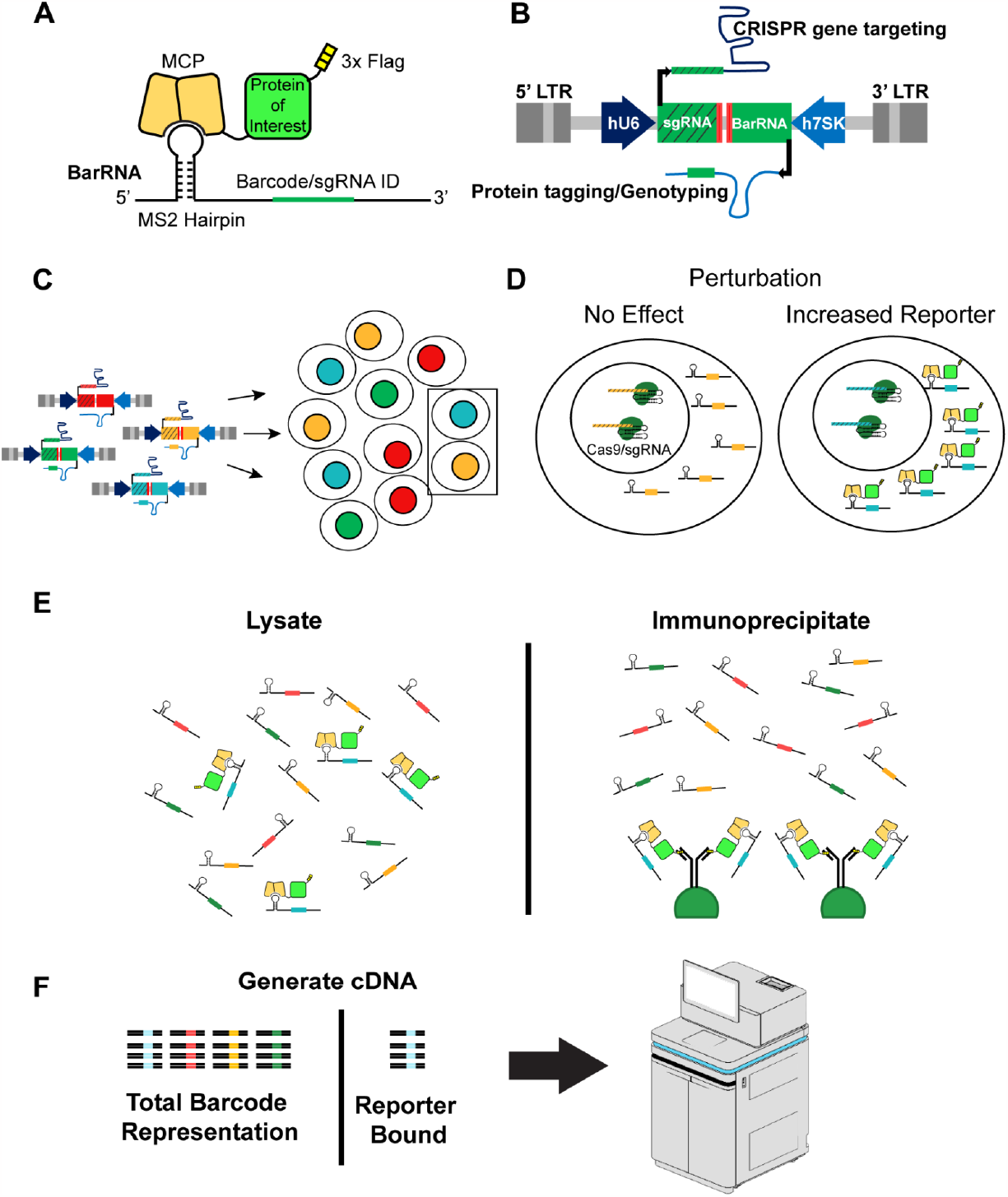
CRISPuRe-Seq by immunoprecipitation. A) Generalized diagram illustrating the core components of a CRISPuRe reporter protein and the BarRNA. B) Diagram of plasmid containing the U6 promoter-driven sgRNA and the 7sk promoter-driven BarRNA. C) Schematic of virally-infected cells containing paired library elements. D) Inset cells from C showing one cell with a perturbation that has no effect on reporter expression and one cell with a perturbation that increases reporter expression. E) Parallel BarRNA purification strategies: Lysate purification represents the total BarRNA population, Immunoprecipitate represents BarRNA bound to the reporter protein. F) BarRNA from both fractions are converted to cDNA and sequenced. The barcode representation in the reporter-bound fraction is then compared to the total barcode representation from the lysate fraction to identify perturbations that alter reporter protein expression.

In the pooled setting, a population of cells is generated where each cell receives a genetic perturbation and expresses its cognate barcode and some amount of a tagged POI (Figure 1C). In the case of protein abundance, perturbations that alter the amount of POI in the cell will alter the amount of POI bound to the corresponding BarRNA (Figure 1D). Relative POI levels are inferred by comparing the distribution of specific BarRNAs after pulldown of the POI to the distribution of BarRNAs in a total RNA fraction from the same cell lysate as measured by high-throughput sequencing (Figure 1E-F). Perturbations that increase the relative amount of POI are enriched for the corresponding BarRNA in the pulldown fraction, while perturbations that decrease the relative amount of POI are depleted for the corresponding BarRNAs.

The POI can be isolated by most standard purification strategies, including immunoprecipitation, co-immunoprecipitation, sedimentation by density and subcellular fractionation. CRISPuRe-seq eliminates the requirement of detecting a fluorescent proxy, does not rely on cell sorting, is highly parallelizable and only uses standard molecular biology techniques. Furthermore, this approach allows for high temporal resolution as cells can be harvested and lysed at any time point. We anticipate that using protein purification as a genetic screening strategy will open the door to many interesting new approaches and discoveries ranging from regulation of protein levels, interaction partners, PTMs and localization.

### Development of an sgRNA/BarRNA co-expression vector

In order to co-express a BarRNA with a corresponding sgRNA, we needed to have both expression cassettes within the same lentiviral construct. BarRNA expression cassettes were cloned into existing lentiviral sgRNA expression plasmids using a convergent transcriptional orientation where the sgRNA and the BarRNA are driven by the strong human U6 and either the 7SK or H1 Pol III promoters respectively (Figure 2A). BarRNA levels are measured by Illumina sequencing after RNA isolation and cDNA synthesis with the incorporation of unique molecular identifiers (UMIs) during reverse transcription. De-duplicated barcode counts are then assigned to their respective sgRNAs in the library (Figure S2A).

**Figure 2.**
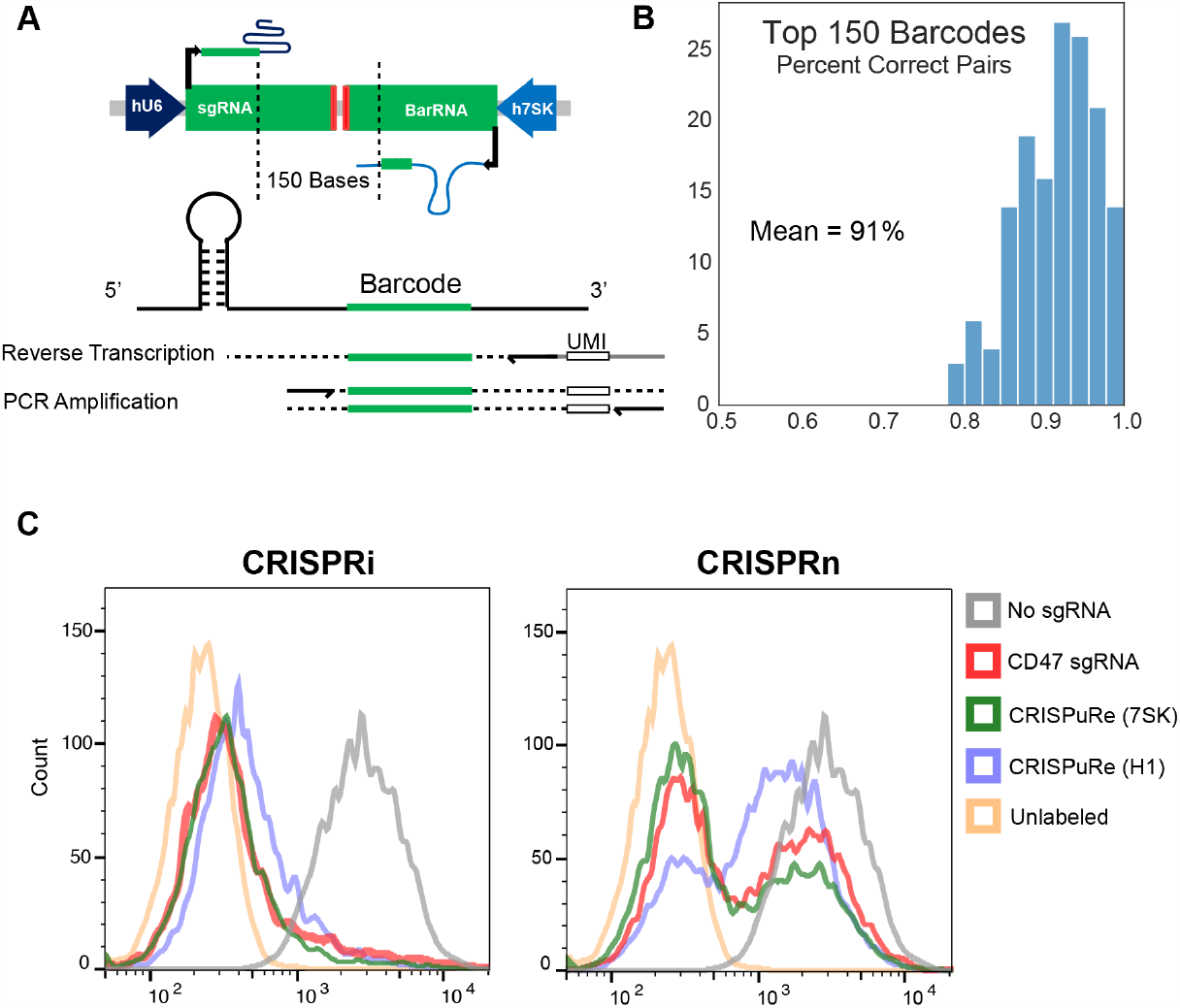
Evaluation of sgRNA and BarRNA co-expression systems. A) The convergent sgRNA/BarRNA arrangement with BarRNA expression driven by the human 7SK Pol III promoter, and the cDNA synthesis and PCR amplification strategy. B) Percentage of correct Barcode/Protospacer pairs for the 150 most abundant barcodes. C) Flow cytometry data showing CRISPR efficacy of 7SK and H1 CRISPuRe constructs. The data for unlabeled cells and for untreated cells (“no sgRNA”) are replicated in each panel as reference.

### Limited recombination between the protospacer and barcode

Recent studies have demonstrated that template-switching during lentiviral integration can result in a ∼6% recombination rate between vector components separated by a common sequence of 96 bases^10^. To minimize recombination between library elements, we cloned the BarRNA and sgRNA expression cassettes in a transcriptionally convergent orientation. This resulted in a 150 bp common sequence between the protospacer of the sgRNA and the 25 nucleotide barcode of the BarRNA.

To estimate the recombination rate using this configuration, we isolated ∼4,000 bacterial colonies from a CRISPuRe cloning reaction using a starting library containing ∼55,000 unique sgRNAs and a random (N)_25_ barcode cassette. Given the number of initial sgRNA in the pool and the diversity of the 25 nucleotide barcode, the vast majority of colonies should represent unique sgRNA/barcode pairs. The plasmid pool was sequenced at 80X coverage and sequences with read counts < 4 were removed to construct the lookup table. This test pool contained 4028 unique sgRNA:BarRNA pairs representing 3537 unique sgRNA protospacers (Figure S2B).

We then packaged the library as a pool to generate lentivirus, infected 500,000 HEK293T cells at a MOI of 0.3 and selected for transduced cells. A PCR amplicon containing the protospacer and barcode was generated from the genomic DNA of infected cells sgRNA:barcode pairs were counted by Illumina sequencing. The coverage per library element at this scale was ∼36x, setting our per-barcode sensitivity to a ∼3% recombination rate.

To estimate lentiviral recombination rates, we considered the top 150 most abundant barcodes (mean read depth of 156) and analyzed their barcode/protospacer pairings. For each barcode the expected reference protospacer pairing represented ∼90% of reads (Figure 2B). A second group of non-reference protospacer pairings with multiple reads represented about 9% of reads (mean of five reads) and a third group of singleton reads.

The recombination rate for each barcode was determined based on the ratio of reads from group 2 vs the reference pairing. The top 150 most abundant barcodes were associated with a mean recombination rate of ∼8.6%, in a mean of 36 transduced cells (Figure S2B). These data suggest that this library configuration likely generates a protospacer/barcode swap <10% of the time, in line with published recombination rates. While small, this barcode swapping represents a source of noise that is inherent to the co-expression system.

### 7SK-CRISPuRe construct is compatible with CRISPRi and CRISPRn

To determine if the convergent arrangement altered sgRNA activity, we performed functional tests in K562 cells stably expressing either KRAB-dCas9 (CRISPRi) or nuclease-active Cas9 (CRISPRn). We expressed either a single sgRNA from the U6 promoter targeting the gene encoding the extracellular membrane protein CD47, or the same sgRNA driven by the U6 promoter in the CRISPuRe configuration with a co-expressed BarRNA driven by either the 7SK or H1 promoter by lentiviral transduction. CD47 expression was measured by fluorescently-labeled anti-CD47 antibody and flow cytometry. The 7SK CRISPuRe construct outperformed the H1 construct in CRISPRi and CRISPR cutting contexts and had the same efficacy as the CD47 sgRNA alone (Figure 2C). Given that the H1 construct showed less efficacy and that the H1 promoter was recently shown to also promote RNA Pol II activity^11^, we selected the 7SK promoter for BarRNA expression.

### Minimal BarRNA exchange during processing

CRISPuRe-seq requires the stable association of the BarRNA with the reporter protein throughout the purification process, ideally with minimal dissociation and RNA exchange. If BarRNA:reporter protein complexes dissociate or freely re-equilibrate after cell lysis, the link between protein phenotype and cellular genotype is broken or scrambled. We chose to develop the system using the coat proteins and cognate RNA hairpins from the MS2 and PP7 bacteriophages based on their well-characterized stability. The reporter proteins consist of an N-terminal tandem dimer of either the bacteriophage MS2-Coat Protein (MCP)^12,13^ or the PP7-Coat Protein (PCP), mClover3^14^, and C-terminal triple-FLAG tag. The MCP domain binds a modified MS2 specific RNA-hairpin sequence with sub-nM affinity and low off-rates compatible with overnight affinity purification protocols (dissociation half-time > 24 hrs at 4 degrees)^15^, and both MCP and PCP have been used in many assays including MS2-TRAP^16^, single-molecule mRNA localization^17^, and RNP tethering^18^.

To evaluate the stability of the MS2-BarRNA with the MCP reporter, we performed a cell mixing experiment and determined the fidelity of BarRNA association with the MCP reporter. HEK293T cells were stably transduced to co-express either 1) a single non-targeting sgRNA:BarRNA pair and tandem MCP or PCP dimer fused to the fluorescent mClover3 and a C-terminal 3X flag (Target Population) or 2) a pool of ∼300 non-targeting sgRNA:BarRNA pairs and tandem MCP or PCP dimer fused to the fluorescent mClover3 without a FLAG tag (Competitor Population) (Figure 3A).

**Figure 3.**
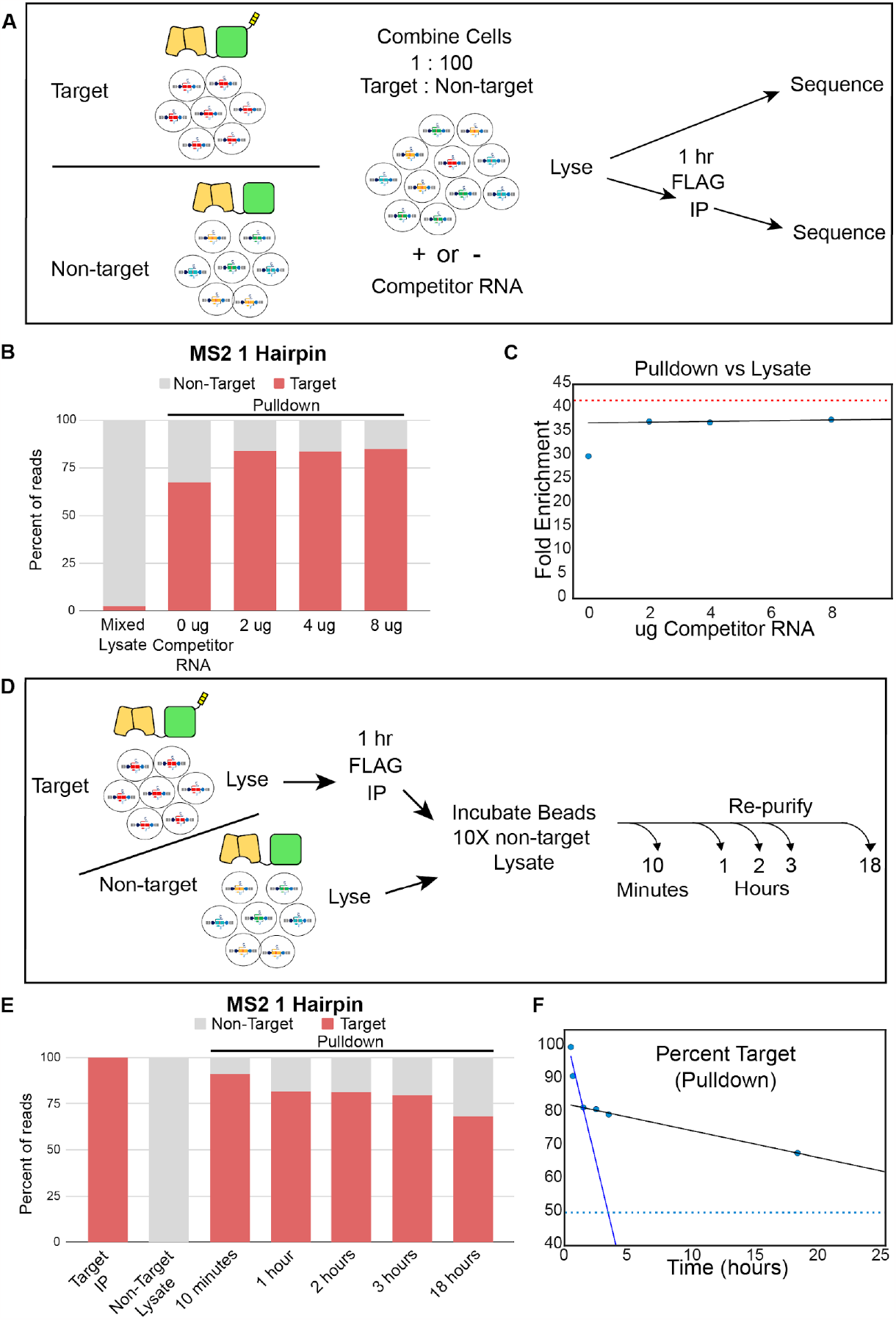
CRISPuRe reporter and BarRNA association is stable. A) Schematic of cell mixing experiment to test Target BarRNA enrichment in HEK 293T cells. B) Graph of Target or Non-target read percentages from either the lysate or the pulldown fractions with or without added competitor RNA. C) Plot of enrichment of Target BarRNA sequence in the IP fraction compared to Lysate fraction. D) Schematic of cell mixing experiment to test exchange rate between bead-bound BarRNA and free BarRNA. E) Graph of Target or Non-target read percentages from the initial FLAG IP, the Non-target lysate, and the pulldown fraction following incubation with Non-target lysate after 10 minutes, 1, 2, 3, and 18 hours. F) Percentage of Target BarRNA reads from pulldown fraction over time.

Cells were mixed at a ratio of 1:100 (Target Population: Competitor Population) and lysed. Ten percent of the lysate was taken for total RNA isolation and a 1 hr immunoprecipitation was performed using anti-Flag magnetic beads on the remaining lysate. Upon purification and sequencing, we found that the IP fraction was highly enriched for the single BarRNA as compared to the lysate fraction (Figure 3B). We hypothesized that BarRNA liberated from the Competitor Population might bind to free protein from the Target Population once the cells were lysed. To test this, we titrated in increasing amounts of a “blocking” BarRNA that contained an MS2 hairpin but was not amplifiable during downstream processing steps. When we included this RNA at a concentration in excess of 2 ug/million cells, which is comparable to the concentration of tRNAs in the lysate, we saw a modest improvement to target RNA enrichment. Inclusion of further RNA beyond this yielded no increase in enrichment (Figure 3C).

As an orthogonal system, many groups have used the similar PCP/PP7 hairpin system. We obtained similar results as with MCP/MS2, but with less efficacy. Initial target BarRNA enrichment was reduced compared to MCP/MS2, though this was mostly rescued by the inclusion of increasing amounts of “blocking” BarRNA (Figure S3A). For this reason, we selected the MCP/MS2 system for RNP barcoding.

Many studies using the MCP-MS2 hairpin system use multiple hairpins to increase MCP binding to an RNA. Addition of a second hairpin to the BarRNA did not improve enrichment compared to a single MS2 hairpin (Figure S3B). Moreover, the inclusion of multiple hairpins in a single BarRNA might facilitate the multimerization of potential MCP-tagged proteins of interest. We therefore proceeded with a single MS2 hairpin in the BarRNA.

### BarRNA binding is sufficiently stable for most purification protocols

To determine how long the reporter protein would remain stably bound to the single-hairpin BarRNA, we performed a variation of the cell mixing strategy (Figure 3D). RNPs from the Target Population lysate were immunoprecipitated, washed, then a 10-fold excess lysate from the Competitor Population was added. We then incubated these samples under IP conditions for increasing amounts of time (10 minutes to 18 hrs) and evaluated pulldown specificity.

We observed a relatively small time-dependent reduction in the relative abundance of Target barcodes in the IP fraction even after 18 hrs of incubation. An initial rapid phase of mixing occurs within the first hour which is followed by a slow linear period of barcode exchange (Figure 3E-F). We suspect this reflects an initial non-specific binding of the added BarRNA/BarRNPs to the beads, followed by slower dissociation of target BarRNA, possibly with exchange with the non-target BarRNA. The slow apparent BarRNA exchange rate that we observe makes 1-2 hour immunoprecipitations highly feasible and might also allow for the possibility of long-duration immunoprecipitation strategies such as tandem purifications.

### CRISPuRe screening identifies positive regulators of Type 1 interferon signaling

Having validated the individual components of CRISPuRe-seq, we next sought to evaluate the approach in a phenotypic screen. As a proof of concept, we performed a set of screens for regulators of type 1 interferon signaling in the Chronic Myelogenous Leukemia cell line, K562. Interferon alpha (IFN-α) activates the canonical JAK/STAT pathway via binding to the receptor proteins IFNAR1 and IFNAR2. This binding activates the Janus Kinases JAK1 and TYK2 which in turn phosphorylate STAT1 and STAT2. A STAT1/STAT2 heterodimer binds to IRF9 to form the ISGF3 (IFN-stimulated gene (ISG) factor 3) complex. This complex translocates to the nucleus and binds to Interferon-stimulated response elements (ISREs) upstream of interferon-stimulated genes to induce their transcription^19^. The negative regulator USP18 can inhibit this cascade by preventing the binding of JAK1 to IFNAR2^20^ (Figure 4A).

**Figure 4.**
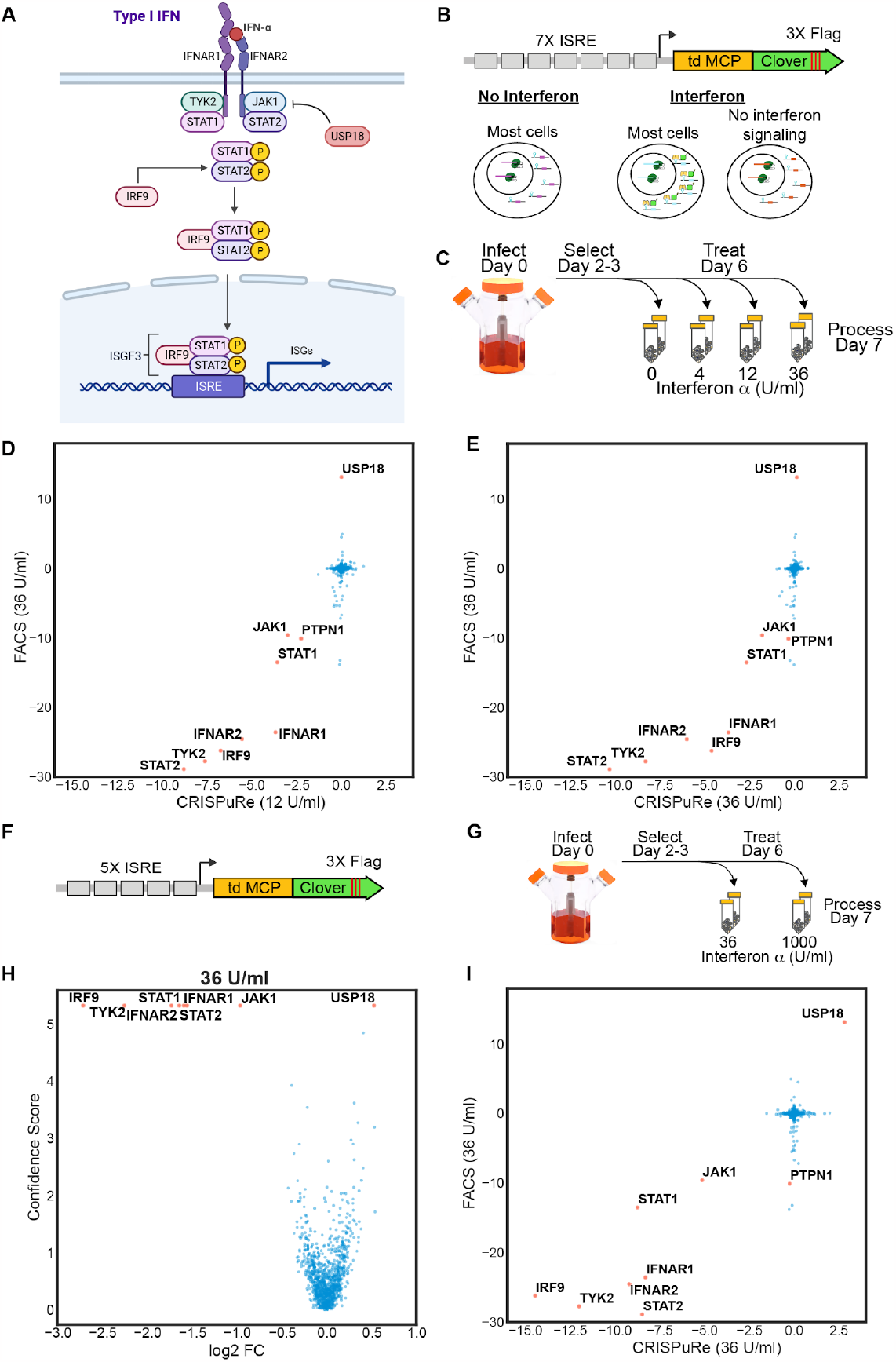
CRISPuRe screening identifies known and possibly new components of IFN alpha signaling cascade. A) Type 1 IFN signaling pathway. B) Schematic of 7X ISRE reporter construct along with a graphic depiction of expected reporter behavior. C) Design and timeline of CRISPuRe dose response screens. D-E) CRISPuRe screen results at increasing Interferon doses (x-axes) plotted against the FACS-based screen performed at 36 U/ml (y-axes). Screen results are plotted as Genescores. F) Schematic of 5X ISRE reporter. G) Design and timeline of 5X ISRE CRISPuRe screens. H) Volcano plot showing result of 5X ISRE reporter screen at 36 U/ml of Interferon alpha. I) The same screen results plotted against the original FACS-based screen at 36 U/ml.

A clonal K562 CRISPRi reporter cell line was generated that expresses the MCP CRISPuRe gene under transcriptional control of a 7X array of ISREs (Figure 4B). We validated the dose responsiveness of the reporter by FACS following a 24-hour treatment with IFN-α, observing a maximal fluorescence response around 1000 U/ml with an EC50 between 12 and 36 U/ml (Figure S4A).

A focused CRISPuRe-BarRNA library of 5930 sgRNA covering approximately 1100 genes involved in inflammation or innate immunity, including the known components of the JAK/STAT pathway was cloned into a lentiviral vector. We then used CRISPuRe to screen this library in the 7X ISRE CRISPuRe reporter cell line at multiple doses of IFN-α (Figure 4C) using Flag immunoprecipitation. In order to benchmark CRISPuRe-seq against standard reporter screening methods, we processed the 36 U/ml arm as a FACS enrichment screen in parallel, collecting the top and bottom 15% of cells based on mClover3 fluorescence.

As expected, CRISPuRe screening in the absence of interferon did not identify any strong candidate regulators (Data not shown). However, screening in the context of 4, 12, or 36 U/ml of IFN-α identified IFNAR1, IFNAR2, JAK1, TYK2, STAT1, STAT2 and IRF9 as strong positive regulators of interferon signaling by CRISPuRe at all doses and by the 36 U/ml FACS screen (Figure 4D-E, S4B). Surprisingly, we identified PTPN1, the gene encoding the PTP1B phosphatase, as a positive regulator of interferon signaling in the context of low dose activation, with a smaller effect at 36 U/ml. While PTP1B has been implicated as a negative regulator of IFN via the promotion of IFNAR1 endocytosis^21^, recent studies have shown that specific PTP1B isoforms derived from alternative splicing can act as positive regulators^22,23^.

### Optimizing reporter dynamic range for identification of positive and negative regulators of interferon signaling

Despite good agreement amongst positive regulators of interferon signaling between FACS and CRISPuRe screens, USP18, a known negative regulator of the pathway, scored only by FACS. We hypothesized that this discordance was due to a sensitivity limit imposed by the BarRNA expression level, such that protein levels in excess of the BarRNA saturate the upper limit of detection.

To test this, we performed a series of mixing experiments using the ISRE reporter and three separate pools of non-targeting BarRNAs. In each experiment one pool received no IFN-α, one pool received 1000 U/ml and the third pool received either 4, 12, or 36 U/ml (Figure S4C). After treatment, cells were mixed at a ratio of 90% untreated, 5% 1000 U/ml and 5% of the variable dose. The mixed cells were lysed and enrichment in the pulldown fraction was determined (Figure S4D). We found significant enrichment at the lowest dose of 4 U/ml, and 36 U/ml was indistinguishable from 1000 U/ml, despite an obvious difference in fluorescence at these doses. This suggested that there was an upper limit to our ability to detect increases in protein expression using our 7X ISRE reporter and that we exceeded that limit somewhere between 12 and 36 U/ml of IFN-α.

In light of this upper sensitivity limit, we generated a lower-gain reporter construct using only 5 ISREs (Figure 4F). We selected a monoclonal line with a dose responsive fluorescence profile but had significantly lower fluorescence than the 7X ISRE construct (Figure S4E). We then ran dose-response screens using the 5X ISRE construct at 36 U/ml and 1000 U/ml (Figure 4G). At 36 U/ml we were now able to identify the previously identified positive regulators as well as the negative regulator USP18 (Figure 4H-I), while at 1000 U/ml we could only identify the positive regulators, consistent with fluorescence levels at which saturation was observed in the 7X reporter (Figure S4A,D-F).

## Discussion

These results demonstrate that pooled CRISPuRe screening by immunoprecipitation allows for increased throughput by virtue of parallel sample processing and no requirement for specialized equipment while generating results comparable to a FACS-based screen. The ability to perform simultaneous screens at different treatment doses allows for a more nuanced interrogation of the biology in question which might lead to the identification of different candidate genes. The throughput of typical reporter screens is FACS-limited, whereas CRISPuRe screens are limited by liquid handling and readily amenable to automation. This is ideal for screens in which multiple treatments from a single reporter cell line are to be tested, allowing for broader interrogation of gene-environment interactions.

We have demonstrated this approach in the context of a transcriptional reporter where we have used the enrichment of BarRNA as a proxy for protein abundance. However, with respect to screening by immunoprecipitation, the functional unit of this approach is not the protein, but the epitope used for protein isolation. Perturbations that alter the abundance of a specific epitope should be screenable with this approach in the absence of any changes in absolute protein levels. Furthermore, this approach should not be limited to immunoprecipitation as a purification strategy as any fractionation method that preserves the integrity of the reporter/BarRNA association should be a screenable phenotype. Thus we anticipate that CRISPuRe will be a broadly useful screening approach that complements existing techniques with unprecedented molecular specificity.

## Supporting information

Scripts for processing raw sequencing data

Plasmid maps

Screen Data

## Data Availability

The data underlying this article are available in the article and in its online supplementary material.

## Acknowledgements

We thank Jacqueline Villalta and John Yong for helpful discussions and comments. We thank Dennis Lin, Bryan King, John Yong, Carmela Sidrauski and Dan Gottschling for reviewing the manuscript and helpful discussions.

