## Supplemental Figures for "Pooled screening for biochemically-defined phenotypes using CRISPuRe-Seq"

### Figure 2 Supplement

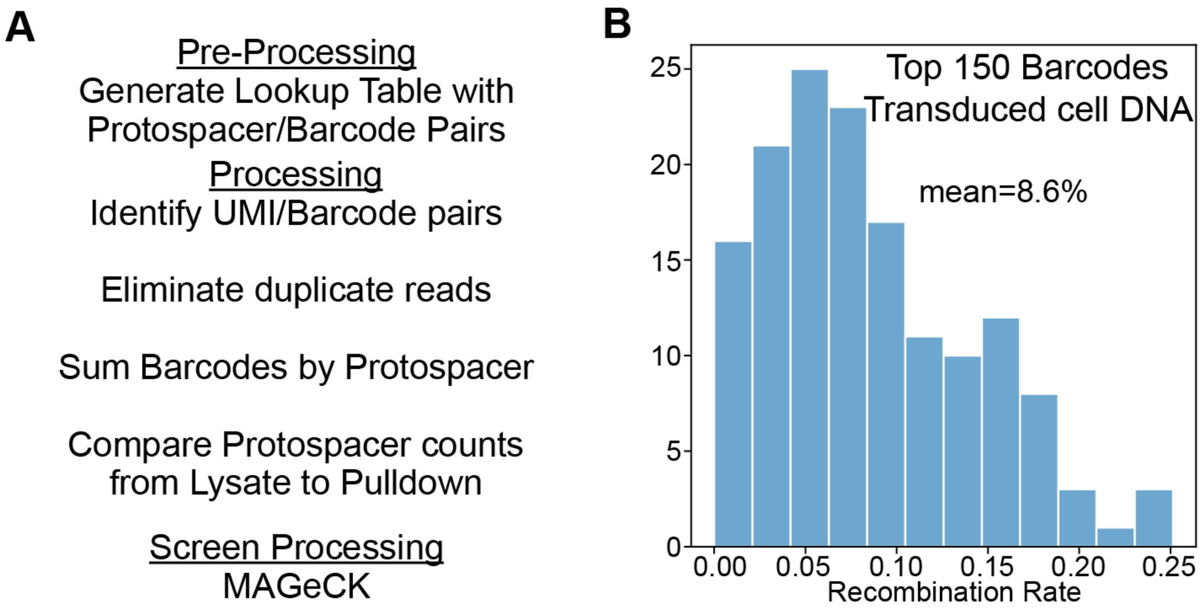


**Supplemental Figure 2. BarRNA Design and CRISPR efficacy.** A) Schematic of the pre-processing and processing pipeline for extracting and tabulating barcode data from sequencing files. B) Recombination rates for the top 150 Barcodes.

### Figure 3 Supplement

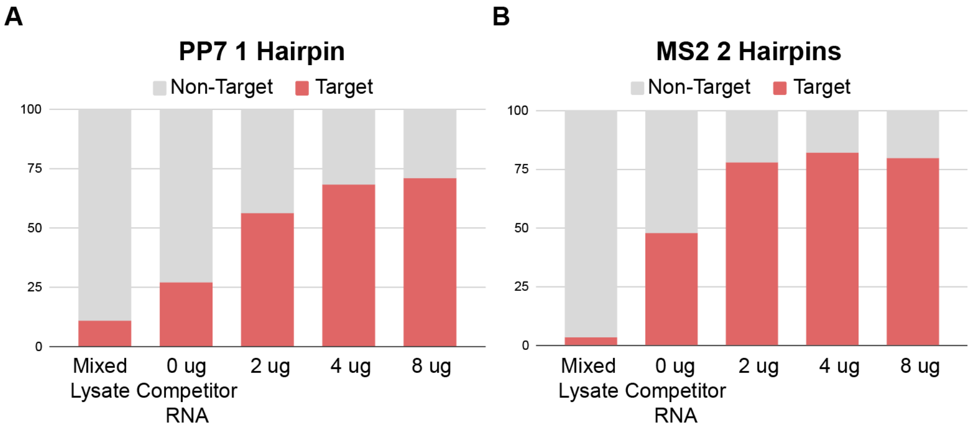


**Supplemental Figure 3. CRISPuRe reporter and BarRNA association is stable.** A) Graph of Target or Non-target read percentages from either the lysate or the pulldown fractions with or without added competitor RNA using the PCP reporter and a BarRNA with one PP7 hairpin . B) Graph of Target or Non-target read percentages from either the lysate or the pulldown fractions with or without added competitor RNA using the MCP reporter and a BarRNA with two MS2 hairpins.

### Figure 4 Supplement

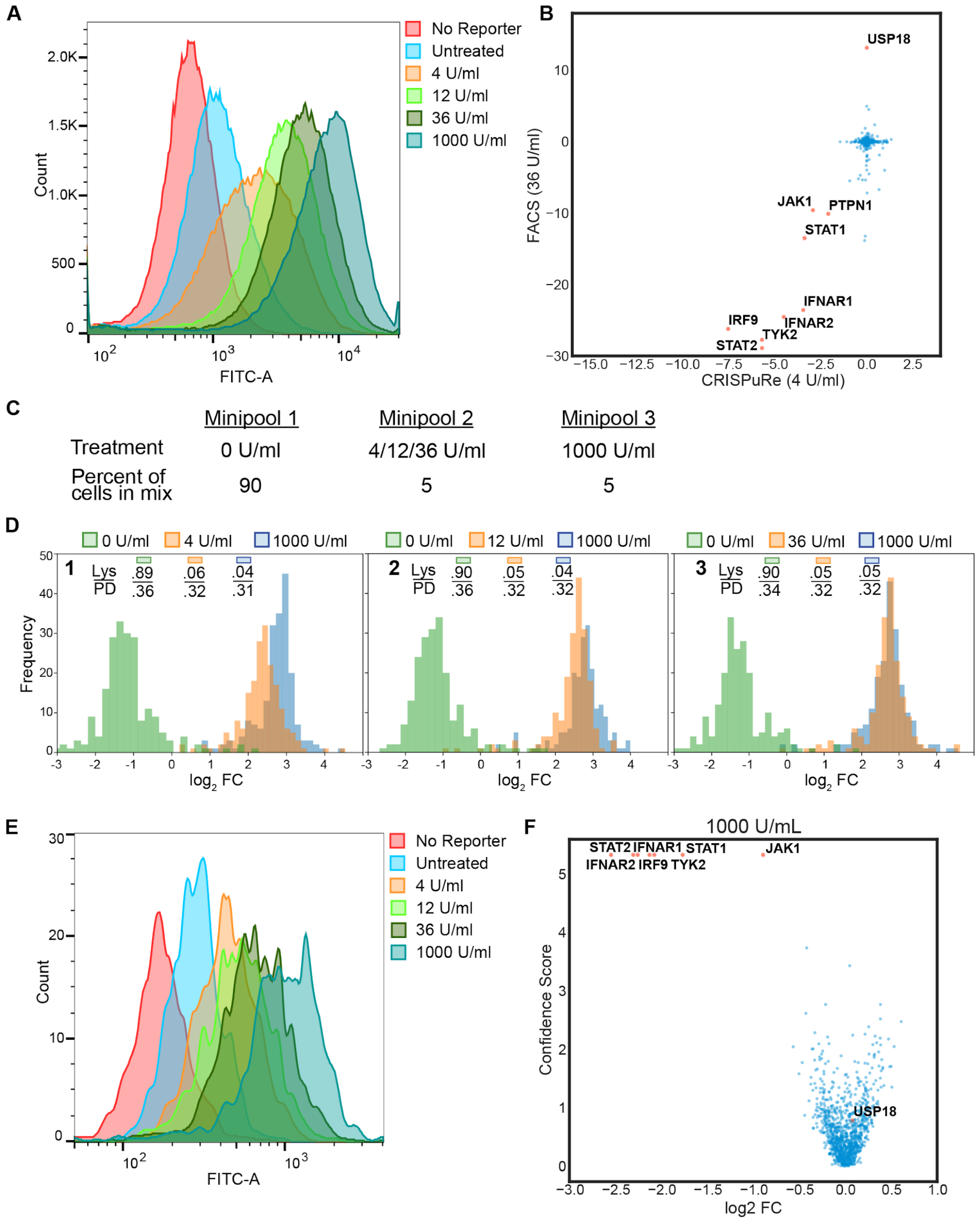

**Supplemental Figure 4. High baseline expression of the CRISPuRe ISRE reporter limits the capacity to detect increases in expression.** A) Flow cytometry data for mClover fluorescence using the 7X ISRE CRISPuRe reporter at increasing doses of interferon alpha. B) 7X ISRE CRISPuRe screen results at 4 U/ml interferon alpha. C) Outline of minipool mixing experiment to test limits of BarRNA enrichment using the 7X ISRE reporter. D) Plots of BarRNA enrichment for three mixing experiments (Panels 1-3). Inset are the percentage of barcode reads from each minipool in the lysate and pulldown fractions for each experiment. E) Flow cytometry data for mClover fluorescence using the 5X ISRE CRISPuRe reporter at increasing doses of interferon alpha. F) 5X ISRE CRISPuRe screen results with an interferon treatment of 1000 U/ml.
