## Supplementary material for "Pooled screening for biochemically-defined phenotypes using CRISPuRe-Seq": Scripts for processing raw sequencing data: CRISPuRe_mageck2.sh

```
#!/bin/bash
#echo "$1"
dir="$PWD/.."
mkdir -p "$dir/$1/Processed"
mkdir -p "$dir/$1/Processed/mageck"
mkdir -p "$dir/$1/Temp"

for i in $2 $3 $4 $5
do 
	filelist=($(find . -name "$i*gz"))
	#echo "$filelist"
	file_1=${filelist[0]}
	file_2=${filelist[1]}
	echo "$file_1"
	echo "$file_2"
	a=$(gzip -cd $file_1 | dd ibs=2048 count=20 | grep -E -no GCAGC[A,G,C,T]{25}CGCT | wc -l)
	b=$(gzip -cd $file_2 | dd ibs=2048 count=20 | grep -E -no GCAGC[A,G,C,T]{25}CGCT | wc -l)
	echo "$a"
	echo "$b"
	if ((a > b))
	then	
		zgrep -E -no GCAGC[A,G,C,T]{25}CGCT $file_1 | awk '{sub(/:/," ",$1)};1'>"$dir/$1/Temp/${i}_bar_lines.txt"
		zgrep -E -no TCTC[A,G,C,T]{15}TGAC $file_2 |awk '{sub(/:/," ",$1)};1'>"$dir/$1/Temp/${i}_umi_lines.txt"
		join <(sort -k 1,1 < "$dir/$1/Temp/${i}_bar_lines.txt") <(sort -k 1,1 < "$dir/$1/Temp/${i}_umi_lines.txt") > "$dir/$1/Processed/${i}_umibars.txt" 
	else
		zgrep -E -no GCAGC[A,G,C,T]{25}CGCT $file_2 | awk '{sub(/:/," ",$1)};1'>"$dir/$1/Temp/${i}_bar_lines.txt"
        	zgrep -E -no TCTC[A,G,C,T]{15}TGAC $file_1 |awk '{sub(/:/," ",$1)};1'>"$dir/$1/Temp/${i}_umi_lines.txt"
        	join <(sort -k 1,1 < "$dir/$1/Temp/${i}_bar_lines.txt") <(sort -k 1,1 < "$dir/$1/Temp/${i}_umi_lines.txt") > "$dir/$1/Processed/${i}_umibars.txt"
	fi

	rm "$dir/$1/Temp/${i}_bar_lines.txt"
	rm "$dir/$1/Temp/${i}_umi_lines.txt"
done


python $HOME/CRISPuRe/CRISPuRe_mageck.py $1 "$dir/$1/Processed/${2}_umibars.txt" "$dir/$1/Processed/${4}_umibars.txt" 1
python $HOME/CRISPuRe/CRISPuRe_mageck.py $1 "$dir/$1/Processed/${3}_umibars.txt" "$dir/$1/Processed/${5}_umibars.txt" 2

python $HOME/CRISPuRe/CRISPuRe_mageck2.py $1 "$dir/$1/Processed/CTable_rep_1.txt" "$dir/$1/Processed/CTable_rep_2.txt"
```
