## Supplementary material for "Pooled screening for biochemically-defined phenotypes using CRISPuRe-Seq": Scripts for processing raw sequencing data: CRISPuRe_mageck2_ISRE.py

```
#!/usr/bin/env python3
import subprocess
import sys
import os


#Import libraries
import pandas as pd

import numpy as np

# Reading in the library of Barcode/Protospacer/Gene pairings
libs = pd.read_csv('/home/dtharris/Data/Experiments/2_CRISPuRe-Seq/CRISPUREILUTs/Lib102020.csv', sep=',', header=0,skipinitialspace=True)
inflam_lib=libs[libs['Sublibrary']=='inflammation/innate immunity']


Dir = sys.argv[2].split('/Processed')[0]

Rep1_ctable = pd.read_csv(sys.argv[2], sep='\t', header=0,skipinitialspace=True)
Rep2_ctable = pd.read_csv(sys.argv[3], sep='\t', header=0,skipinitialspace=True)

Rep1_ctable.columns=['sgRNA','Gene','one_1.initial','two_1.final']
Rep2_ctable.columns=['sgRNA','Gene','one_2.initial','two_2.final']

Rep1_2_merged = pd.merge(Rep1_ctable,Rep2_ctable, on = ['sgRNA','Gene'],how= 'outer')
Rep1_2_merged=Rep1_2_merged[['sgRNA','Gene','two_1.final','two_2.final','one_1.initial','one_2.initial']]

Rep1_2_merged=Rep1_2_merged[Rep1_2_merged['Gene']!='negative_control']
Rep1_2_merged=Rep1_2_merged[Rep1_2_merged['Gene'].isin(inflam_lib['Gene'])]

Rep1_2_merged['two_1.final'][np.isnan(Rep1_2_merged['two_1.final'])]=0
Rep1_2_merged['two_2.final'][np.isnan(Rep1_2_merged['two_2.final'])]=0
Rep1_2_merged['one_1.initial'][np.isnan(Rep1_2_merged['one_1.initial'])]=0
Rep1_2_merged['one_2.initial'][np.isnan(Rep1_2_merged['one_2.initial'])]=0


Rep1_2_merged=Rep1_2_merged[~Rep1_2_merged['Gene'].str.contains('non-target')]

Rep1_2_merged.to_csv(Dir +'/Processed/'+sys.argv[1]+'_paired_CTable.txt',sep = '\t',header=True, index=False)

mageckcall= 'mageck test -k '+Dir +'/Processed/'+sys.argv[1]+'_paired_CTable.txt -t two_1.final,two_2.final -c one_1.initial,one_2.initial -n '+Dir+'/Processed/mageck/'+sys.argv[1]+' --sort-criteria "pos" --gene-lfc-method "median" --paired'
os.system(mageckcall)
```
