## Supplementary material for "Pooled screening for biochemically-defined phenotypes using CRISPuRe-Seq": Scripts for processing raw sequencing data: CRISPuRe_mageck.py

```
#!/usr/bin/env python3
import subprocess
import sys
import pandas as pd
import numpy as np

Dir = sys.argv[2].split('/Processed')[0]
Dir_umi = Dir + "/Processed/"
Expt = sys.argv[1]
Samp_one = sys.argv[2]
Samp_two = sys.argv[3]
rep=sys.argv[4]

Output = Dir + "/Processed/CTable_rep_"+rep+".txt"

#Read in 7sk library file and Sample one umibars file
a = pd.read_csv('/home/dtharris/Data/Experiments/2_CRISPuRe-Seq/CRISPUREILUTs/Lib102020.csv', sep=',', header=0,skipinitialspace=True)
b = pd.read_csv(Samp_one, sep=' ', header=None,skipinitialspace=True)

#Isolate barcodes from umibars file
b.columns = ['Line','Barcode','UMI']
b = b.drop('Line', axis = 1)
b['Barcode']=[s[5:30] for s in b['Barcode']]

#Merge library file with umibars
one_lib = pd.merge(b,a, on = 'Barcode')

#Deleting b since it is merged into one_lib
#del a
del b

#Grouping by UMI and Barcode to look for duplicates
one_lib_grouped = one_lib.groupby(['UMI','Barcode','Protospacer','Gene','Sublibrary']).size().reset_index()

#Sorting values in ascending order to calculate gini coefficient for library construction
one_lib_sorted = one_lib_grouped.sort_values(by = 0, ascending=True)

#Cleaning house
del one_lib
del one_lib_grouped

# Eliminate duplicates
one_lib_uni = one_lib_sorted.drop_duplicates(subset=['Barcode','UMI','Gene'],keep = 'first')

one_lib_uni=one_lib_uni.drop([0],axis=1)

#Cleaning house
del one_lib_sorted

#Counting the number of unique reads of each barcode
one_lib_uni_grouped=one_lib_uni.groupby(['Barcode','Protospacer','Gene','Sublibrary'],as_index=False).size().reset_index()

#Cleaning house
del one_lib_uni

#Preparing dataframe for merge with sample two data
one_lib_uni_grouped=one_lib_uni_grouped.drop(['index'],axis=1)
one_lib_uni_grouped.columns=['Barcode','Protospacer','Gene','Sublibrary','one_count']
one_lib_proto=one_lib_uni_grouped.groupby(['Protospacer','Gene','Sublibrary'])['one_count'].sum().reset_index()

del one_lib_uni_grouped

d = pd.read_csv(Samp_two, sep=' ', header=None,skipinitialspace=True)

#Copy of all of the steps performed above, this time on sample two data
d.columns = ['Line','Barcode','UMI']
d = d.drop('Line', axis = 1)
d['Barcode']=[s[5:30] for s in d['Barcode']]
#print(d.head(5))
two_lib = pd.merge(d,a, on = 'Barcode')
#pull_totalbars = d[~d['Barcode'].isin(a['Barcode'])]
del d
#del a
two_lib_grouped = two_lib.groupby(['UMI','Barcode','Protospacer','Gene','Sublibrary']).size().reset_index()

del two_lib

two_lib_sorted = two_lib_grouped.sort_values(by = 0, ascending=True)

del two_lib_grouped

two_lib_uni = two_lib_sorted.drop_duplicates(subset=['Barcode','UMI','Gene'],keep = 'first')

del two_lib_sorted
#Calculate number of unique UMI-Barcode pairings in sample two

two_lib_uni=two_lib_uni.drop([0],axis=1)
two_lib_uni_grouped=two_lib_uni.groupby(['Barcode','Protospacer','Gene','Sublibrary'],as_index=False).size().reset_index()
del two_lib_uni
two_lib_uni_grouped=two_lib_uni_grouped.drop(['index'],axis=1)

two_lib_uni_grouped.columns=['Barcode','Protospacer','Gene','Sublibrary','two_count']

two_lib_proto=two_lib_uni_grouped.groupby(['Protospacer','Gene','Sublibrary'])['two_count'].sum().reset_index()

del two_lib_uni_grouped

#Merging sample one data with the sample two data
Final_merge=pd.merge(one_lib_proto,two_lib_proto, on=['Protospacer','Gene','Sublibrary'],how='outer')
#print(Final_merge.shape)
#For those Barcodes that are not present in sample two that get converted into NaN, this makes them a 0
Final_merge.fillna(0,inplace=True)
#print(Final_merge.head(5))

Final_merge=Final_merge[Final_merge['one_count']>3]

CTable = "_CTable_rec.txt"
#This code generates the final count table for processing in Mageck
count_table = Final_merge[['Protospacer','Gene','one_count','two_count']].copy()
count_table.columns = ['sgRNA','Gene','one.initial','two.final']
count_table.loc[:,'two.final']=count_table['two.final'].astype(int)
count_table.to_csv(Output,sep = '\t',header=True, index=False)
```
